# Limited protection against early-life cytomegalovirus infection results from deficiency of cytotoxic CD8 T cells

**DOI:** 10.1101/2024.07.10.602923

**Authors:** Luís Fonseca Brito, Eleonore Ostermann, Anna Perez, Silvia Tödter, Sanamjeet Virdi, Daniela Indenbirken, Laura Glau, Anna Gieras, Renke Brixel, Ramon Arens, Adam Grundhoff, Petra Arck, Anke Diemert, Eva Tolosa, Wolfram Brune, Felix Rolf Stahl

## Abstract

Differential antiviral T cell immunity in early life impacts the clinical outcome of Cytomegalovirus (CMV) infection, but the underlying mechanisms are not well understood. Here, we found delayed enrichment of early-life murine CMV-specific CD8 T cells due to a general deficiency of αβ T cells. Adoptive transfer of naïve adult T cells into neonates did not protect due to a blockade of CD8 but not of CD4 effector T cell differentiation. Early-life deficiency of critical signal 3 cytokines during T cell priming resulted in the appearance of non-cytotoxic CD8 effector T cells whereas the effector phase of adult-primed T cells was not disrupted in neonates. Accordingly, we found an overall low number of antiviral human CD8 T cells in newborns with congenital CMV. Together, this study suggests defective CD8 T cell immunity as an important factor explaining the higher risk for CMV disease in the early-life phase.

## Introduction

Early-life Cytomegalovirus (CMV) infection, in humans and mice, is characterized by prolonged viral replication and increased risk of disease^1–4^. While CMV is generally considered to be an opportunistic pathogen causing predominantly mild or asymptomatic infection in immunocompetent hosts^5^, congenital CMV (cCMV) or infection of preterm low-birth-weight newborns is the most important infectious cause of permanent disabilities in infants^6^. To date, there is no protective vaccine available and therapeutic options are limited, not curative, and bear the risk of severe adverse effects^7^. Accordingly, cCMV causes significant health burden^8^ and urgently requires investigation of disease pathophysiology.

CMV induces exceptionally strong T cell responses that are crucial for virus control and regularly lead to ∼15 % of the circulating CD8 T cell pool to be specific for CMV epitopes^9^. Thus, it is reasonable to assume that age-related differences in anti-CMV T cell immunity are responsible for the increased vulnerability in the early life. Indeed, neonates exhibit an altered composition of peripheral blood T cell compartments and differential antigen-recognition patterns compared to adults^10^. However, virus-specific effector T cells are present in newborns after infection with human CMV (HCMV) ^11^ and in neonatal mice challenged with murine CMV (MCMV) ^3,12,13^, indicating that in general, the very young host can acquire anti-CMV T cell immunity. Nevertheless, the presence of these effector T cells does not preclude CMV disease, and it is currently not known to which extent these early-life effector T cells exhibit protective functions.

Here, we delineate age-related differences in anti-CMV T cell immunity. In mice, we demonstrate that the expansion of endogenous MCMV-specific T cells early in life is delayed due to an overall low frequency of peripheral T cells. Adoptive transfers of adult T cells into neonates generated CD8 effector T cells that were low in cytotoxicity, and this correlated with their non-protective function in the young host. These non-cytotoxic CD8 T cells resulted from altered early-life T cell priming conditions, whereas the T cell effector phase itself was not disrupted. Consistent with this, we confirmed an overall lower CD8 T cell response in infants with cCMV indicating that the early-life susceptibility to viral infection is linked to dampened cytotoxic T lymphocyte (CTL) induction.

## Results

### Delayed expansion of MCMV-specific CD8 T cells in the early life

To investigate the pathophysiology of the high early-life risk for CMV disease we infected neonatal and adult mice by administering MCMV in the respiratory tract^3^. Hypothesizing that T cell immunity is involved in the delayed MCMV control in the lungs of neonates (Fig 1A) ^3^, we tracked MCMV-specific T cells in peripheral blood using pMHC-I tetramer staining. We detected CD8 T cells recognizing MCMV immunodominant epitopes M38, M45, and m139^14^ after seven days post-infection (dpi) in adults (Fig 1B+C). In neonates, MCMV-specific CD8 T cells could not be detected before 10 dpi and the frequency of these cells was lower (Fig 1B+C). We also analysed the accumulation of M25-specific CD4 T cells^15^ and detected more cells stained with an MHC-II tetramer in infected than in control animals in both, adults and neonates (Fig S1A+B). To follow up on the low numbers of MCMV-specific CD8 T cells in neonates we assessed the presence of lymphocytes in spleen of non-infected animals during the very early postnatal life period. At the day of birth, we found populations of B220^+^ B and CD3^-^NK1.1^+^ natural killer (NK) cells, but very few CD3^+^ T cells (Fig 1D+E and S1C). Within the first days of life, the microanatomy of the spleen underwent a distinct organization, and the numbers of CD3^+^ T cells expressing TCRβ and CD4 or CD8β in the white pulp increased (Fig 1D+E). The relative number of B cells also increased, albeit to a lesser extent than T cells, whereas the frequency of NK cells remained rather stable within the first four weeks of life. By calculating the absolute number of lymphocytes relative to the animal body weight we found an approximately 10.000-fold increase of T cells in the spleen within the first week of life (Fig 1F). Taken together, we concluded that the low precursor frequency of T cells in the early life could explain the delayed enrichment of MCMV-specific CD8 T cells and reduced control of lung infection.

**Fig 1.**
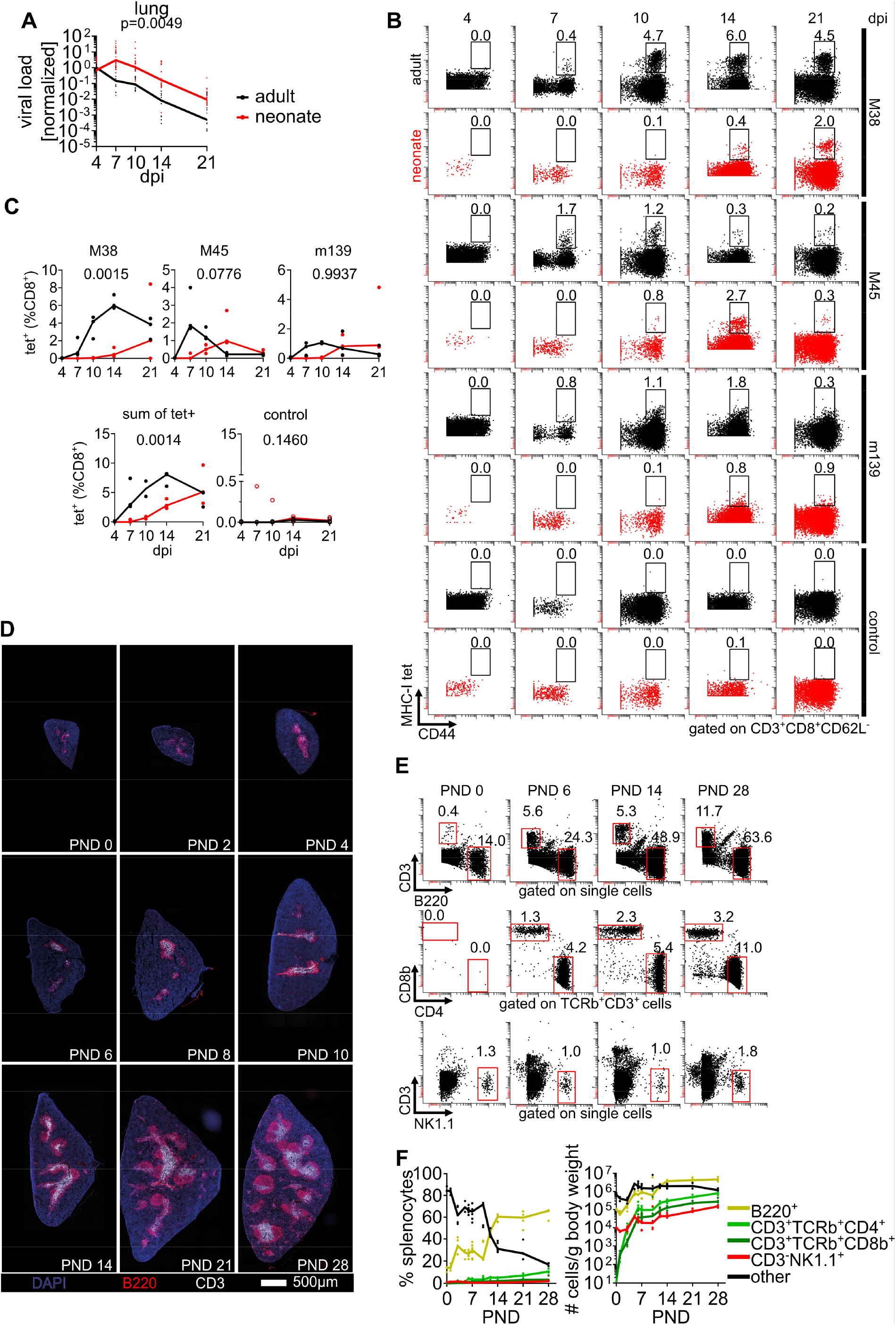
Delayed enrichment of MCMV-specific CD8 T cells in early life (A-C) Neonatal and adult mice were infected with MCMV and analysed at indicated days post infection (dpi). (A) Lung viral loads of adult and neonatal mice after MCMV-infection. Data normalized to the median value at 4 dpi are displayed. (B+C) Representative flow cytometry plots and (C) pooled analysis of frequency of peripheral blood CD8 T cells stained for MCMV peptide MHC-I tetramers as indicated. Non-infected mice were stained simultaneously with a pool of all three MHC-I tetramers as controls. (D-F) Non-infected mice were analysed for appearance of major lymphocyte populations. (D) Immunohistology of mouse spleens stained as indicated at post-natal days (PND) as indicated. (E) Representative flow cytometry plots of major lymphocyte populations in the spleen. (F) Frequency (left panel) and absolute number of cells normalized to animal body weight (right panel) of major lymphocyte populations in spleen. Data were acquired from more than two independent experiments (A, n= 8-29 per time point; B and C, n=3 per time point; D - F, n=3-7 per time point). Lines in (C) and (F) connect median value of each time-point. Statistical difference between frequency of MCMV-specific CD8 T cells in (C) was calculated via a 2-way ANOVA and the p value is provided above each graph.

## Naïve adult T cells adoptively transferred into MCMV-infected neonates do not protect from infection

To test whether low numbers of T cells accounted for the delayed control of MCMV infection in early life, we adoptively transferred purified polyclonal CD3^+^ T cells obtained from secondary lymphoid organs of adult act-eGFP mice into neonates (Fig 2A). Transferred cells homed to lymphoid organs and led to an approximately 60-fold increase of T cells in spleens within two days post intraperitoneal injection, reaching similar cell per body weight numbers as found in four-week-old mice (Fig 2B+C).

**Fig 2.**
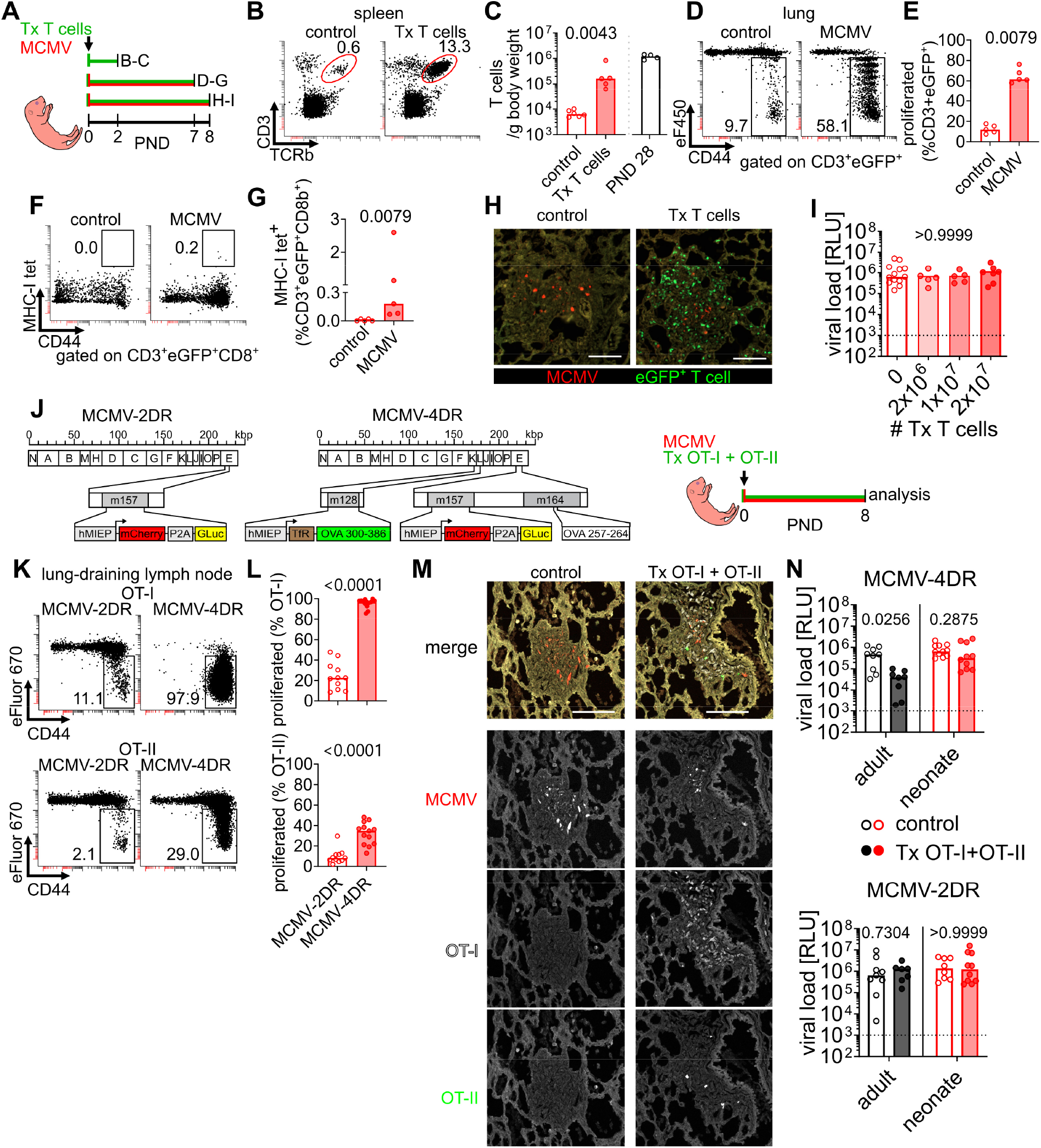
Adoptive transfer of adult naïve T cells into neonates is not protective against MCMV (A) Experimental setup for (B-I): (B+C) 2x10^7^ purified T cells obtained from adult act-eGFP transgenic mice (eGFP) were transferred (Tx) into neonatal mice and analysed after 2 days. (D-G) 10^7^ eGFP^+^ T cells were stained with cell proliferation dye eF450 and adoptively transferred into neonatal mice which were simultaneously infected with MCMV, control mice received cells but were not infected. Animals were analysed at 7 days post cell transfer. (H+I) Various numbers of eGFP^+^ T cells were adoptively transferred into neonatal mice, which were simultaneously infected with MCMV, control mice received no T cells. Animals were analysed at 8 days post infection (dpi). GLuc, *Gaussia* luciferase; hMIEP, human major immediate early promotor; OVA, chicken ovalbumin; TfR, transferrin receptor sequence. (B) Representative flow cytometry plots of spleen T cells. (C) Absolute numbers of T cells calculated to animal body weight 2 days after adoptive transfer. Cell numbers of non-treated PND 28 animals are depicted from Fig 1F for comparison. (D) Representative flow cytometry plots of eGFP^+^ T cells isolated from lungs indicating eF450 signal loss with each proliferation cycle. (E) Quantitative analysis of proliferated eGFP^+^ T cells isolated from lungs. (F) Representative flow cytometry plots of eGFP^+^ CD8^+^ T cells isolated from lungs simultaneously stained with M38-, M45-, and m139-pMHC tetramers (G) Quantitative analysis of MCMV-specific tetramer^+^ eGFP^+^ CD8^+^ T cells isolated from lungs. (H) Immunohistology of lungs indicating localisation of transferred eGFP+ T cells in nodular inflammatory foci (NIF) (I) Lung viral loads after adoptive transfer of various numbers of naïve T cells. RLU, relative light units. (J) Experimental setup for (K-N): MCMV-2DR contains no additional inserted sequences besides the fluorophore and luciferase reporters whereas MCMV-4DR encodes for sequences of chicken ovalbumin. Neonatal wildtype or adult *Rag2^-/-^Il2rg^-/-^* mice were infected (10^4^ or 2x10^5^ PFU, respectively) with either MCMV-2DR or MCMV-4DR and received adoptive transfers of 5x10^6^ OT-IeCFP and 5x10^6^ OT-IIeGFP proliferation dye-labelled cells, control animals did not receive T cells. Animals were analysed at 8 dpi. (K) Representative flow cytometry plots of T cells isolated from lung draining lymph nodes indicating eF450 signal loss with each proliferation cycle. (L) Quantitative analysis of proliferated T cells. (M) Immunohistology of MCMV-4DR -infected neonatal lungs indicating localisation of transferred OT-IeCFP and OT-IIeGFP T cells next to MCMV-infected cells in NIF. (N) Lung viral loads after infection with MCMV-4DR (upper panel) or MCMV-2DR (lower panel) and adoptive transfer of OT-I and OT-II cells. Data were acquired from two or more experiments (B and C, n=5-6; D-G, n=5; H-I, n=5-13, K-N, n=7-11 animals per group). Scale bars, 100 µm. Statistical differences between groups were calculated with Mann-Whitney U (C, E, G, L and N) or Kruskal-Wallis (I) tests the p values are provided above each graph.

Next, we combined adoptive T cell transfers with MCMV infection and found the adult T cells to acquire CD44 membrane expression and proliferate in neonatal mice in response to virus challenge (Fig 2D+E). Moreover, a fraction of the CD8 T cells was MCMV-specific, indicating that the increase of T cell precursors indeed augmented the numbers of MCMV-specific CD44^+^ effector T cells at 8 dpi (Fig 2F+G). MCMV causes the formation of nodular inflammatory foci (NIF) in the lung, in which T cells recognize antigens and participate in the control of infection^3,16^. Accordingly, adoptively transferred T cells accumulated in neonatal NIFs (Fig 2H). However, adult T cells did not decrease the number of MCMV-infected cells in neonatal lung NIFs (FigS2A), and even transfer of high numbers of T cells did not reduce the viral load in the lungs (Fig 2I).

To further increase the number of antigen-specific T cells in neonatal NIFs we infected the animals with MCMV-4DR, a recombinant virus encoding for epitopes recognized by OT-I and OT-II transgenic T cells, or with MCMV-2DR as a control (Fig 2J). MCMV-4DR infection of neonatal mice led to significantly higher proliferation of adoptively transferred naïve OT-I and OT-II cells than in mice infected with MCMV-2DR (Fig 2K+L). We found OT-I and, to a lower degree, OT-II cells in NIFs at 8 dpi (Fig 2 M). As a control, we used adult *Rag2*^-/-^*Il2rg*^-/-^ mice which lack major lymphocyte populations. These mice were infected with either of the two MCMV recombinants, and received the same numbers of naïve OT-I and OT-II cells as neonates. Importantly, this equals to a 20-fold lower number of adoptively transferred T cells into adults calculated per gram bodyweight. While antigen-specific T cells reduced viral loads in MCMV-4DR-infected *Rag2*^-/-^*Il2rg*^-/-^ adult mice, there was no significant effect in neonatal mice or animals infected with MCMV-2DR (Fig 2N and S2B). Adoptive transfer of polyclonal T cells into adult *Rag2*^-/-^ mice decreased viral loads when approximately 100 T cells were present in NIFs^17^. Here, although up to 250 OT-I T cells were found in neonatal NIFs, these cells failed to significantly reduce the number of infected cells or inflamed tissue (Fig S2C-E). The findings highlight that adoptive transfer of adult naïve T cells into neonates can compensate for the deficiency of this cell population in the early-life period, lead to antigen-specific proliferation, gain of effector marker expression, homing into sites of MCMV infection, but has no influence on virus load in the lung.

## Early-life T cell priming impedes differentiation into antiviral cytotoxic T lymphocytes

Since transferred adult naïve T cells were unable to protect from early-life MCMV infection we hypothesized that age-related factors modulate T cell differentiation and effector function. Therefore, we compared the phenotype of approximately 6000 adult T cells after priming in either adult or neonatal mice by single-cell transcriptome profiling (scRNA-seq) (Fig 3A +S3A). T cells isolated from the lungs of MCMV-infected adults and neonates exhibited a distinct transcriptome when compared to non-infected controls (Fig 3B). Expression of *Cd4* or *Cd8* was equally distributed across the four different conditions (FigS3A) and positioned cells into two main clusters (Fig 3C). Most of the cells obtained from the MCMV-infected mice had low *Ccr7* expression indicating that they lost their naïve phenotype (Fig 3C). In line, we identified a higher frequency of T cells with shared T cell receptors (TCRs) after MCMV infection than in control animals, supporting the idea of antigen-dependent proliferation and emergence of effector T cells (Fig 3D). T cells obtained from non-infected neonatal and adult mice had similar RNA expression profiles whereas in cells obtained from MCMV-infected mice a notable age-related difference was evident (Fig 3E). To gain insight into CD4 T cell differences *Cd4*^+^*Cd8a*^-^ cell clusters were annotated according to their gene expression signatures into naïve (*Tcf7*, *Ccr7*, and *Sell*) and cycling (*Mki67* and *Pclaf*) T cells, regulatory T cells (Treg) (*Foxp3* and *Il2ra*), and T-helper 1 cells (Th1) (*Cd44*, *Ly6c2*, and *Cxcr6*) ^18,19^(Fig 3F+G). All CD4 T cell subpopulations were found in the four different conditions (Fig 3H). MCMV infection decreased the relative number of naïve T cells, which on the other side led to a higher frequency of cycling T cells and Th1 cells (Fig 3H). Tregs were found in similar numbers independently of age or infection (Fig 3H + S3B). However, there was a trend that adult T cells primed in MCMV-infected neonates were more likely to differentiate into Th1 cells (Fig 3H) and these cells showed a distinct gene expression pattern (Fig 3I, table S1). There was higher expression of pro-inflammatory, anti-apoptotic, and cell cycle-related genes in neonates (e.g. *Ifng*, *Birc5, Pclaf*) (Fig 3I). Moreover, CD4 T cells obtained from neonates had a higher cytotoxicity module score in the MCMV-infected groups (Fig 3J), but not in the controls (FigS3C). Priming in MCMV-infected neonates led to a stronger clonal T cell response than in adults (Fig 3K), but with comparable relative distribution into the four different CD4 subpopulations (Fig 3L). Taken together, there were age-related differences in CD4 T cell priming with a stronger Th1 response in cells obtained from neonates.

**Fig 3.**
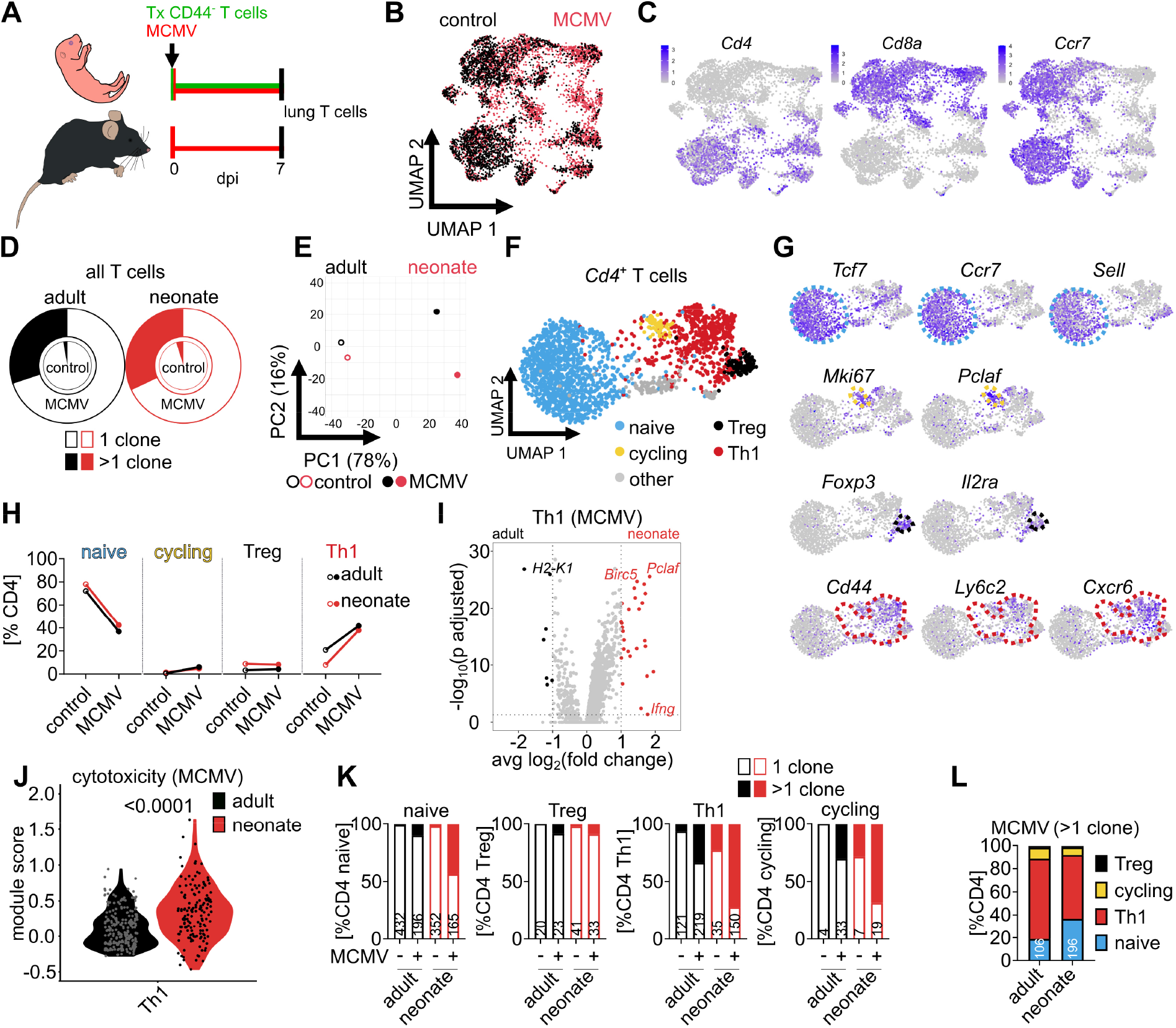
CD4 T cells primed in neonates acquire a Th1-like phenotype with cytotoxic potential (A) Experimental setup for Fig 3 + 4: Neonatal (n=3) and adult (n=3) mice were infected with 10^4^ or 2x10^5^ PFU MCMV, respectively. 10^7^ purified CD3^+^CD44^-^ naïve T cells obtained from adult eGFP mice were transferred into neonatal mice. Control animals (n=3 neonates and n=3 adults) were not infected with MCMV. After 7 days, adoptively transferred eGFP^+^ lung T cells in neonates and endogenous lung T cells from adults were isolated, and pooled for single-cell transcriptome profiling (scRNAseq) combined with TCR sequencing. (B) Unsupervised RNAseq analysis and UMAP dimensionality reduction of all cellsalgorithm, colours indicate infection status. (C) Cell expression of genes as indicated. (D) Relative distribution of T cells with identical TCRs indicating clonal enrichment, filled area represent clonal cells. (E) Principal component analysis for the four different groups. (F) Subanalysis and UMAP dimensionality reduction of *Cd4*^+^*Cd8a*^-^ T cells. (G) Classification of CD4 T cells into subpopulations by expression of characteristic genes as illustrated. (H) Frequency of the four groups assigned to the different CD4 T cell subpopulations. (I+J) (I) Differential gene expression and (J) cytotoxicity module score of Th1 cells obtained from MCMV-infected adults and neonates. (K) Relative distribution of the clonal (filled section of bars) and not clonal (open bars) CD4 T cell subpopulations of each group with numbers indicating the absolute cell counts. (L) Relative distribution of clonally expanded CD4 T cells into subpopulations. Data acquired by one experiment with n=3 animals per group. Statistical difference in (J) was calculated with Mann-Whitney U test and the p value is provided above the graph.

Next, we annotated the *Cd8*^+^*Cd4*^-^ CD8 T cell clusters into naïve (*Sell*, *Tcf7*, *Ccr7*), cycling (*Mki67*, *Pclaf*), and effector T cells (*Gzmb*, *Cd44*, *Ccl5*) ^20,21^(Fig 4A+B). MCMV infection led to a decrease of naïve and an increase of cycling and effector CD8 T cells in both, adults and neonates (Fig 4C). Comparison of effector T cells revealed differential gene expression with differentiation-promoting genes *Zeb2* and *Id2* and cytotoxicity-related genes such as *Ccl5*, *Gzmk*, and *Gzma* being higher expressed in adults, whereas CD62L-encoding gene *Sell* and differentiation-repressing genes *Tcf7*, *Lef1,* and *Zeb1* were more abundant in neonates (Fig 4D, table S2). Accordingly, T cells isolated from neonates and adults were differently distributed within the effector T cell cluster (Fig 4E). Thus, we performed a sub-cluster analysis of CD8 T effector cells and named the according subpopulations as Teff 1 (high in e.g. *Cd44*, *Ly6c2*, *Gzmm* but also *Sell*), Teff 2 (high in e.g. *Ifit1*, *Ifit3*, and *Gzmb*), Teff 3 (high in e.g. *Cxcr6*, *Cxcr3*, and *Ccl5*) and Teff 4 (high in e.g. *Ccl5*, *Gzma*, *Cx3cr1*, *Zeb2*, and *Tbx21*) (Fig 4F+G, table S2). All Teff1-4 subpopulations were present in neonates and adults (Fig 4H). However, T cells isolated from non-infected neonates clustered more as Teff1 and less as Teff4. In response to MCMV infection changes in Teff1 and Teff2 were comparable between neonates and adults but there were fewer cells in neonates that acquired a Teff3 and Teff4 phenotype (Fig 4H). Accordingly, we found a higher frequency of lung CD8 T effector cells to be positive for membrane protein staining of CXCR6 and intracellular CCL5 in MCMV-infected adults than in neonates (Fig S4A-C). Next, module score analysis revealed that in adults Teff2-4 but not Teff1 exhibited high cytotoxicity function in response to MCMV infection indicating that true CTLs are found within clusters Teff2-4 (Fig 4I + FigS4D). In contrast, neonatal Teff2 - the CD8 effector subpopulation that strongly increased in response to neonatal MCMV (Fig 4H) - had a low cytotoxicity module score (Fig 4I). Moreover, differentiation into cytotoxic Teff3 and Teff4 was minor in neonates when compared to adults (Fig4H) explaining the overall lower expression of cytotoxicity-related genes in CD8 effector T cells in neonates (Fig4D). Furthermore, clonal expansion in response to MCMV as an indicator for antigen-specific CD8 T cell response was higher in adults (Fig 4J). In neonates, clonally expanded CD8 T cells remained in a cycling state whereas in adults most CD8 T cells acquired an effector phenotype (Fig 4K+L). Taken together, there was a significant age-related difference in the CD8 T cell effector response with lower clonal enrichment and cytotoxicity in MCMV-infected neonates than in adults, resulting in significantly fewer antiviral CTLs in the early life.

**Fig 4.**
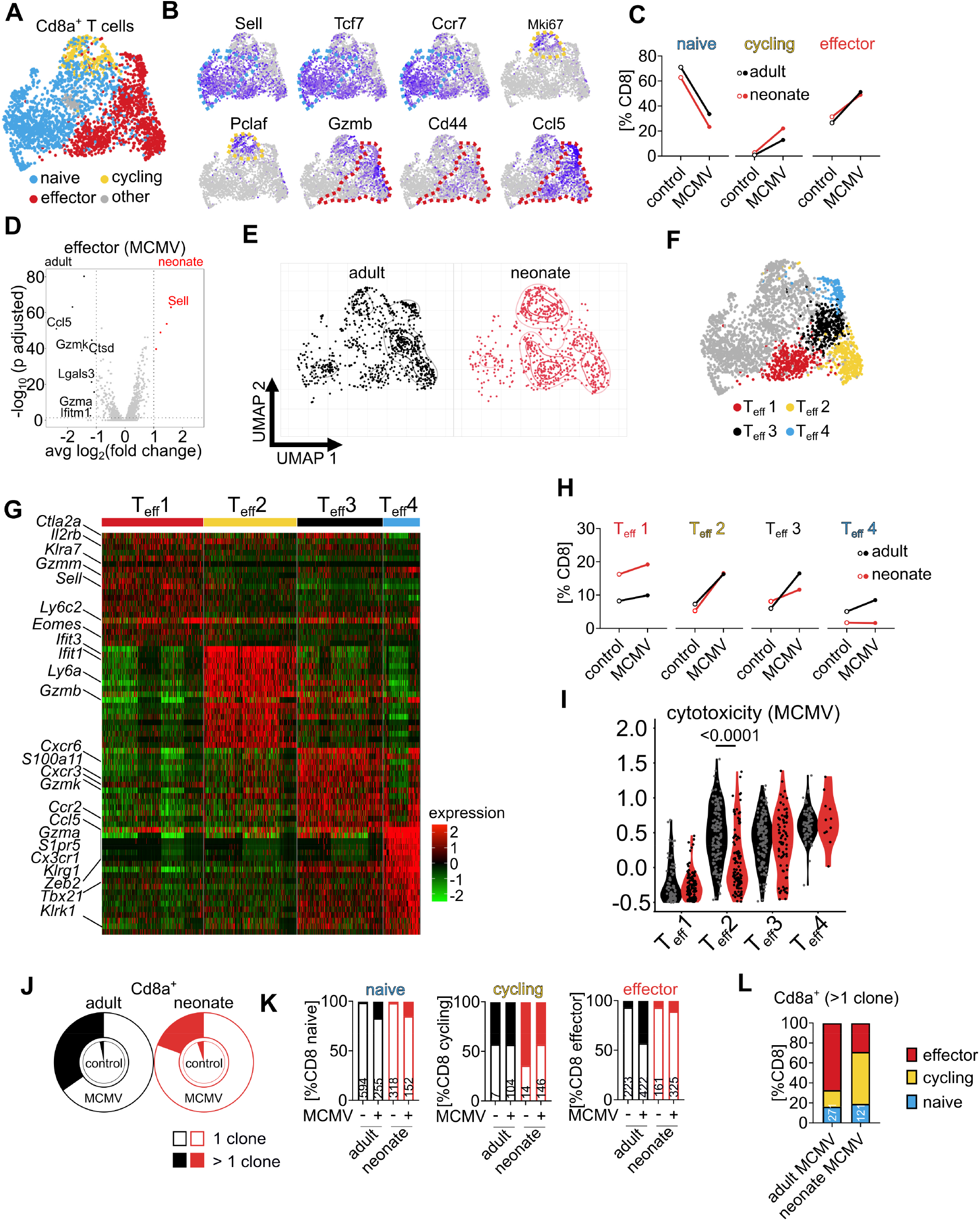
CD8 T cells primed in neonates do not acquire an antiviral CTL phenotype (A) Subanalysis and UMAP dimensionality reduction of *Cd8a*^+^*Cd4*^-^ T cells. (B) Classification of CD8 T cells into subpopulations by expression of characteristic genes as illustrated. (C) Frequency of the four groups assigned to the different CD8 T cell subpopulations. (D) Differential gene expression of effector CD8 T cells isolated from infected neonates and adults. (E) UMAP dimensionality reduction of CD8 T cells isolated from MCMV-infected animals. (F) Subclassification of effector CD8 T cells clusters. (G) Heatmap of top 20 highly expressed signature genes in CD8 T cell effector subclusters. (H) Frequency of the four groups assigned to the different CD8 effector T cell subpopulations. (I) Cytotoxicity module score of CD8 effector T cell subpopulations from MCMV-infected animals. (J) Relative distribution of CD8 T cells with identical TCRs indicating clonal enrichment, filled area represent clonal cells. (K) Relative distribution of the clonal (filled section of bars) and not clonal (open bars) CD8 T cell subpopulations of each group with numbers indicating the absolute cell counts. (L) Relative distribution of clonally expanded CD8 T cells into subpopulations. Data acquired by one experiment with n=3 animals per group. Statistical difference in (I) was calculated with Mann-Whitney U test and the p value is provided above the graph.

## Ineffective early-life CD8 T cell immunity is associated with alterations in cytokine signaling conditions

The effector phenotype of antiviral CD8 T cells is highly influenced by the inflammatory cytokine milieu within the infected tissue^22^. Thus, we quantified cytokines known to be involved in CD8 T cell priming in lungs in steady state or in MCMV-mediated inflammatory conditions at 5 dpi. In general, more cytokines could be detected in non-infected adults than in neonates (Fig5A-L). MCMV infection induced a shift in the local adult cytokine environment to higher concentrations of Interleukin-2 (IL-2), IL-7, and IL-15 which was not detected in neonates (Fig 5A-D). Moreover, there was a robust age-related difference for IL-27, IL-33, and Interferon-α (IFN-α) which were found in higher concentrations in adults (Fig5E-G). In contrast, there was a trend for more IL-10, IL-1β, and IFN-β (Fig5H-J) in neonates and IFN-γ as well as IL-6 (Fig5K+L) were significantly higher in inflamed neonatal lungs. Thus, there were remarkable age-related differences in signal 3 cytokines that likely impact T cell priming and effector differentiation.

**Fig 5.**
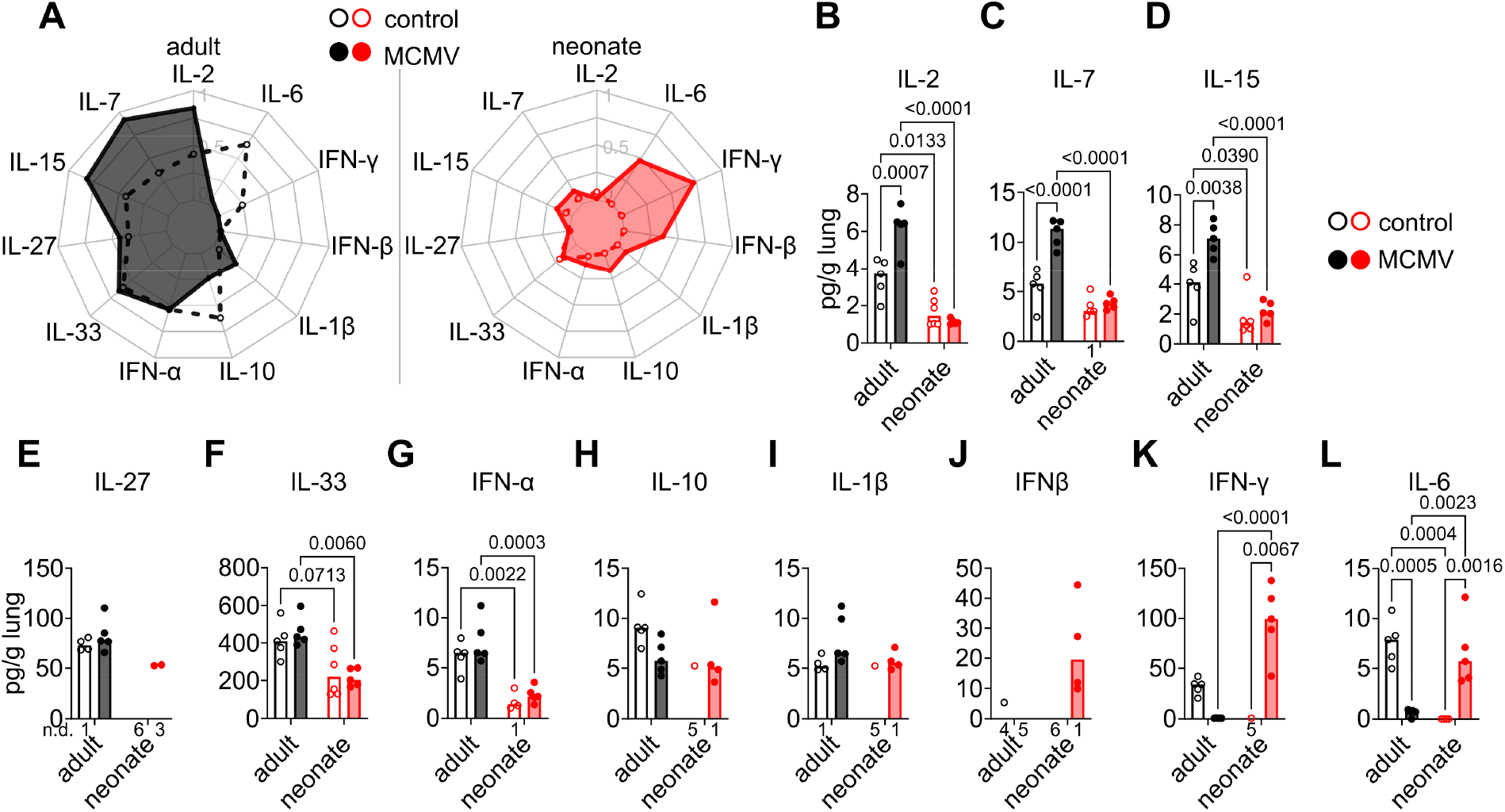
Alterations in priming conditions in the early-life phase (A-L) Relative concentration of cytokines in steady state and MCMV-infected lungs at 5 dpi as indicated. Data acquired from two (E-P) independent experiments. Values in A depict a normalized scale. Statistical differences were calculated with 2-way ANOVAs and the p values are provided above the graphs.

## MCMV-primed adult effector T cells protect from early-life MCMV lung infection

To confirm that T cells primed in adults are more protective against MCMV we isolated CD44^+^ T cells from infected adult lungs and adoptively transferred these into neonatal mice (Fig 6A). At 8 dpi we found 1.9-fold more T cells in lungs of mice that received adult effector T cells as compared to mice without cell transfer (Fig 6B+C). CD44 expression was higher on T cells in mice which received the adoptive transfers indicating that these cells had maintained their effector phenotype (Fig 6D+E). Adoptive transfer of effector T cells reduced inflammation in lungs (Fig 6F), defined by a reduced number and size of NIFs (Fig 6F-H), and the number of MCMV-infected cells within NIFs (Fig 6F+I). In line, the overall virus load was lower in lungs of mice that received the T cell treatment (Fig 6J). Accordingly, the higher cytotoxicity score found in adult-primed CD8 T effector cells correlated with protective function and these cells could reduce MCMV infection in neonatal mice.

**Fig 6.**
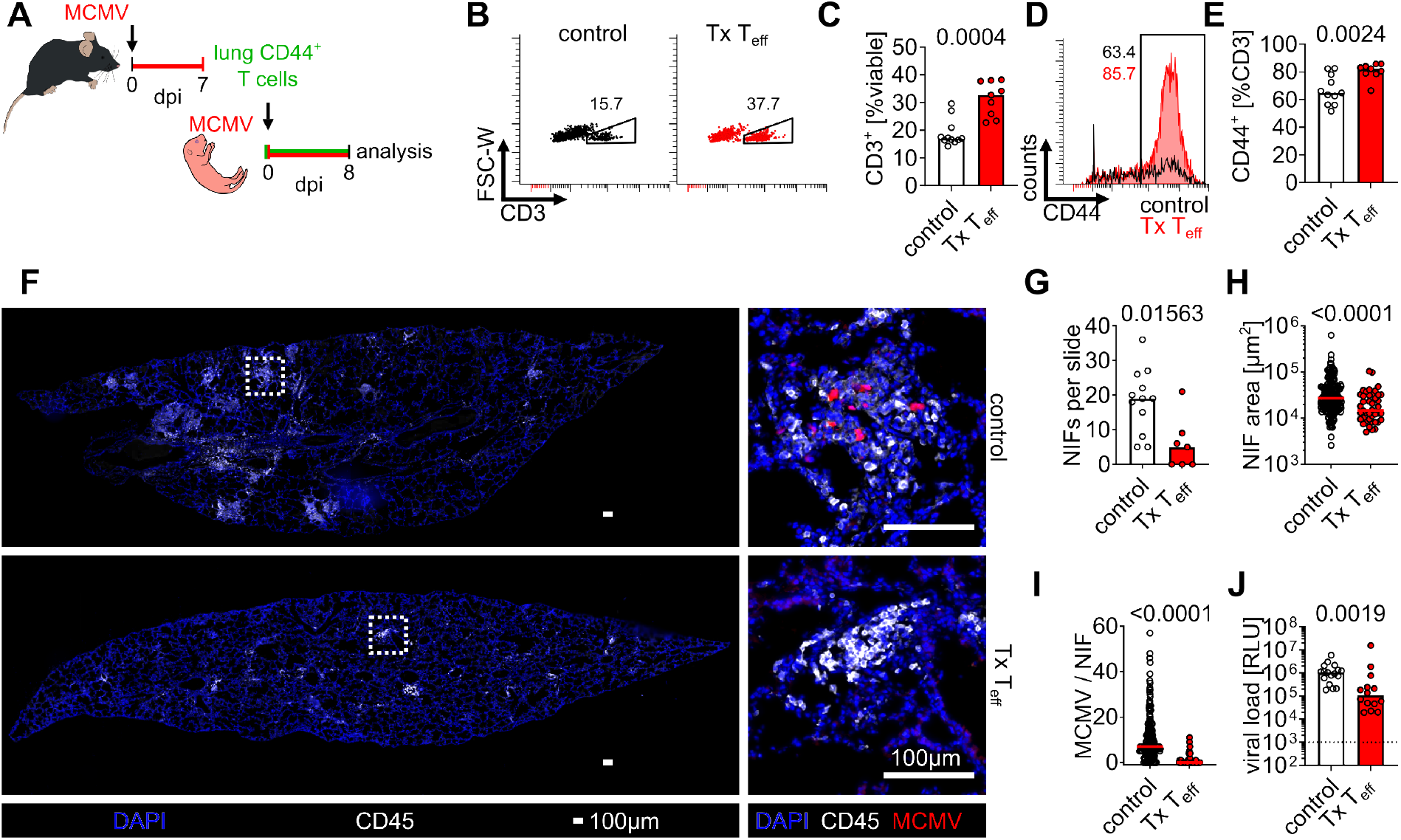
Effector T cells primed in adults protect from early-life MCMV lung infection (A) Experimental setup for (B-J): CD3^+^CD44^+^ effector T cells were purified from adult lungs 7 dpi and 8x10^6^ cells per animal were transferred into MCMV infected neonatal mice which were then analysed 8 dpi, controls did not receive T cells. (B and C) (B) Representative flow cytometry plots and (C) quantitative analysis of CD45^+^ cells obtained from neonatal lungs at 8 dpi. (D and E) (D) Representative flow cytometry histograms and (E) quantitative analysis of CD44 expression in isolated lung T cells. (F) Immunohistology of lungs in general view (left panels) and high resolution (right panels) illustrating inflammation and presence of NIFs with MCMV-infected cells. Dotted squares indicate the zoomed areas shown in the right panels. (G-H) Quantitative analysis of immunohistology with (G) number of NIFs per lung slice, (H) NIF area and (I) number of MCMV-infected cells per NIFs. (J) Quantitative analysis of lung viral loads as indicated. Data acquired from four independent experiments. Statistical differences were calculated in C, E and G-J with Mann-Whitney U test and the p values are provided above the graphs.

## Differential T cell immunity in response to congenital HCMV infection

Finally, we investigated if the age-related differences found in the anti-MCMV immune response could be confirmed in individuals infected with HCMV. Thus, we performed multiparameter flow cytometry-based immune phenotyping to screen for differences in PBMC subpopulations in newborns delivered by mothers with maternal HCMV infection. These newborns are at risk for congenital infection and in this cohort 4 out of 23 (17.4 %) were HCMV positive. Of 88 leukocyte subpopulations analysed in newborns, we found differences between HCMV^+^ and HCMV^-^ individuals mainly in the T cell compartment (FigS5A, table S3). Thus, we included additional samples obtained from adult blood donors with known HCMV serostatus to determine age-related differences in lymphocyte composition. We focused on 44 peripheral blood T cell traits (Fig 7A). In general, frequencies of newborn peripheral blood T cell subpopulations differed significantly to those found in adults (Fig 7A+B). Thus, we independently analysed newborn and adult measurements to determine HCMV-associated changes in the T cell compartment. Indeed, samples obtained from HCMV^+^ and HCMV^-^ individuals clustered apart in both, newborns and adults (Fig 7C+D). There were significantly different frequencies of 7 and 15 of adult and newborn T cell subpopulations, respectively (Fig 7E+F). In adults HCMV infection led to changes in the CD4 and CD8 T cell compartment (Fig 7F). In neonates, there were HCMV-dependent differences in several αβ CD4 and CD8 T cells (Fig 7F, G, H, I J) but also in γδ T cells (Fig 7F). Interestingly, there were overlapping HCMV-dependent differences in three T cell subpopulations in adults and newborns, and these were all of the CD8 T cell compartment (Fig 7E+F). In general, we found more CCR7^+^CD45RA^+^ naïve and accordingly less CCR7^-^CD45RA^-^ effector memory (EM) and CCR7^-^CD45RA^+^ terminally differentiated effector memory (TEMRA) CD8 T cells in newborns than in adults (Fig 7K + L, M, N). HCMV infection was associated with a higher frequency of effector CD8 T cells although the relative number of CD8 EM and TEMRA cells remained significantly lower in newborns (aFig 7M+N). CD57^+^ CD8 T cells are known to expand in HCMV-infected individuals^23^ and may assist as a surrogate to quantify HCMV-specific CD8 T cells. The frequency of CD57^+^ CD8 T cells were found in higher frequencies in HCMV-positive individuals than in seronegative controls but composed only a minor fraction in newborns (Fig 7O+P). In summary, HCMV infection had a significant impact on the CD8 T cell compartment in adults and newborns. However, early-life CD8 T cells remained predominantly in a naïve status and there was a lower frequency of virus-induced effector T cells.

**Fig 7.**
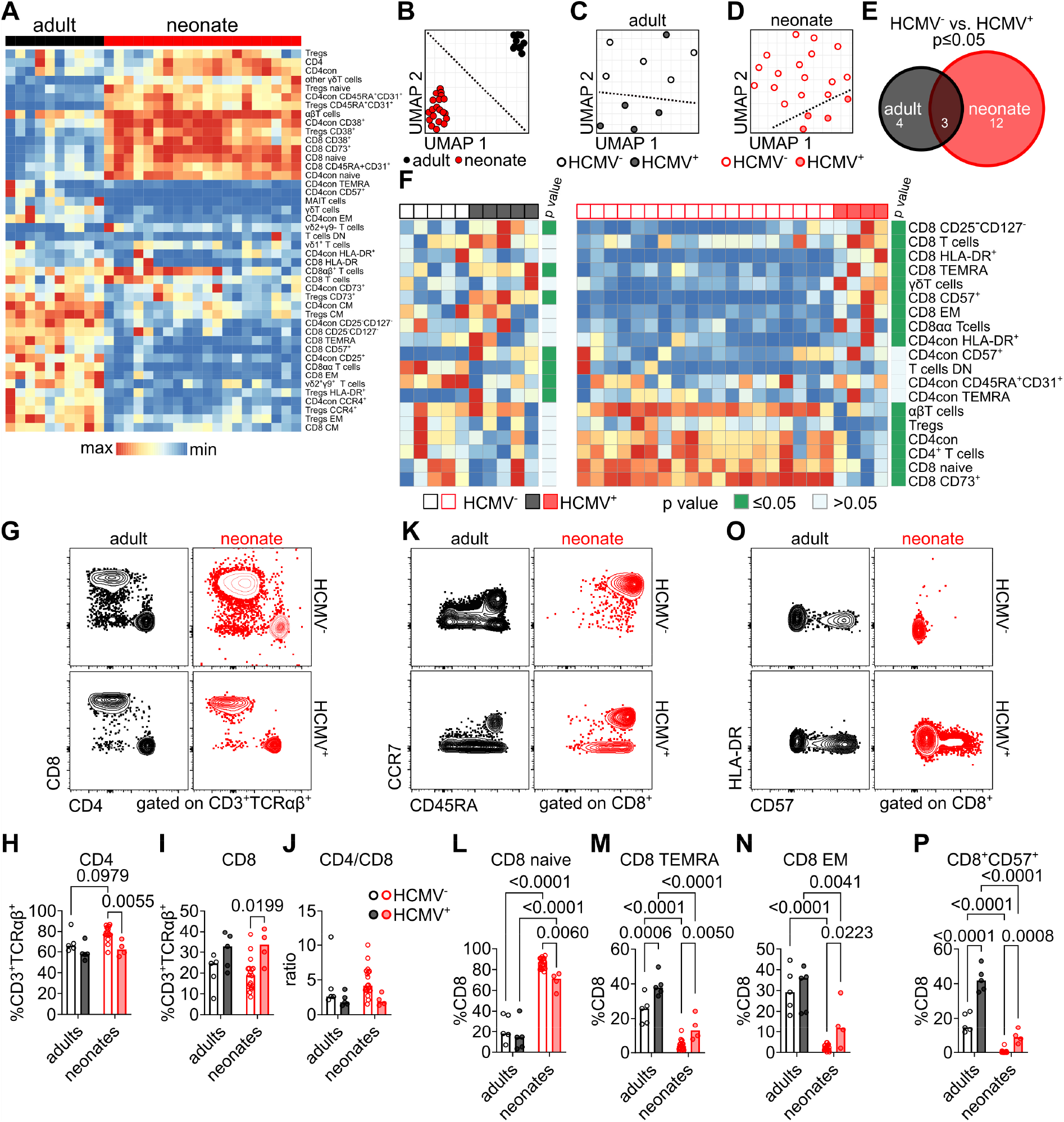
T cell immunity in response to congenital HCMV infection (A) Normalized frequencies of T cell subpopulations in adults and neonates. (B) UMAP dimensionality reduction of T cell subpopulations of adult and neonatal samples. (C) UMAP dimensionality reduction of T cell subpopulations of all adult samples showing HCMV serostatus. (D) UMAP dimensionality reduction of T cell subpopulations of all neonatal samples showing HCMV serostatus. Dotted lines were manually added. (E) Numbers of T cell populations with significantly different frequency depending on HCMV serostatus. (F) Normalized frequencies of the 19 T cell subpopulations that were significantly different depending on HCMV serostatus in adults or neonates. (G -J) (G) Representative flow cytometry plots and (H-J) quantitative analysis of CD4 and CD8 T cells. (K-N) (K) Representative flow cytometry plots and (L-N) quantitative analysis of CD8 T cell differentiation status. (O and P) (O) Representative flow cytometry plots and (P) quantitative analysis of CD57^+^CD8 T cells. Statistical differences were calculated for (E) and (F) by Mann-Whitney U test and for (H-J), (L-N), and (P) by 2-way ANOVA, the p values are provided above each graph. The following abbreviations are used in the figure: con, conventional; TEMRA, terminally differentiated effector memory; MAIT, mucosal-associated invariant T; EM, effector memory; DN, double-negative; CM, central memory.

## Discussion

Upon viral infection, CD8 T cell differentiation is shaped by multiple stimuli during the priming phase including TCR peptide binding, co-stimulation, and cytokine signaling. Integration of these stimuli drives the expression of several transcription factors, chemokine receptors and effector molecules that affect the generation and the effector function of antiviral CTLs. In this study, we found distinct priming conditions to lead to significant differences in the phenotype and function of virus-specific CTLs in early life.

Murine respiratory tract *Cd44*-, *Gzmb*-, and *Ccl5*-expressing CD8 effector T cells encompassed four subpopulations that differed in their expression of effector molecules. Among these Teff3 and Teff4 were high in expression of *Gzmk*, *Gzma*, *Cx3cr1*, *S1pr5*, *Klrg1*, *Cxcr6*, and *Zeb2* arguing for their late differentiation state^24^. Together with Teff2 these three subpopulations exhibited a high cytotoxicity score upon adult MCMV challenge and are likely predominantly involved in control of lung infection. In contrast, Teff1 exhibited signatures of less differentiated effector T cells and had a low cytotoxicity score. With these criteria approximately 40 % of CD8 T cells primed in adults could be classified as CTL with a high cytotoxicity score and potential for anti-MCMV effector function *in situ*. This was contrasted by the fact that CD8 T cells primed in early life were less likely to acquire a Teff3 or Teff4 phenotype, and those cells that clustered as Teff2 exhibited a low cytotoxicity score, resulting in an overall CTL-like phenotype of only ∼10 % of CD8 T cells. Interestingly, in HCMV^+^ individuals we found a median frequency of 42 % and 9 % of CD57^+^ CD8 T cells in adults and newborns, respectively. These cells are known to expand upon HCMV infection^23^, express cytolytic molecules^25^, exhibit partial coexpression of NKG2C and likely include terminally differentiated cells^26^. Similarly, there are more CD8 EM and CD8 TEMRA cells in HCMV^+^ adults than in newborns, the latter being associated with better control of HCMV infection in transplant recipients^27^. Moreover, although fetal CD8 T cells produce IFN-γ upon peptide stimulation^11^, the amount of cytokines produced after mitogen stimulation was lower than in adults^28^. Thus, both in mice and humans, adults are equipped with higher frequencies of cytotoxic CTLs upon CMV infection indicating their superior capacity to control this pathogen. Accordingly, effector CD8 T cells isolated from adults efficiently reduced MCMV loads in neonates and *ex vivo* expanded donor-derived HCMV-specific CTLs were successfully used for the treatment of HCMV infection in paediatric recipients^29^ arguing that the CD8 T cell effector phase is not disrupted in the young host. Instead, CD8 T cell priming in the early life impairs their differentiation program into cytotoxic CTLs and increases their vulnerability to infection.

We found significantly higher production of IL-2, IL-7, and IL-15 in MCMV-infected adult lungs whereas these cytokines were low in neonatal mice and did not increase in concentration in response to infection. Importantly, these three cytokines belong to the common receptor γ-chain (γc) family and are notorious for their role in T cell proliferation, homeostasis, and differentiation^30^. IL-2 is known to drive CD8 effector T cell function and expression of cytotoxic molecules^31^. Moreover, protection by adoptive transfer of CD8 effector T cells into MCMV-infected irradiated adult mice was increased when combined with the application of recombinant IL-2^32^. IL-7 is a critical survival factor for T cells and important for generating CD8 T cell memory^33^. *IL-15ra*^-/-^ mice are deficient in CD8 T cells^34^ and IL-15 is critical for the maintenance of effector memory CD8 T cells in infected lungs after MCMV infection^35^. Furthermore, stimulation of CD8 T cells with IL-15 *ex vivo* upregulates gene expression of *Ccl5* and *Cxcr6*^36^ and here, we found CD8 T cells to exhibit higher expression of these genes and proteins in adults. A previous study suggested that CCL5 itself may recruit CD8 T cells in MCMV-infected lungs^37^ and CXCR6 was shown to position CTLs next to IL-15 trans-presenting dendritic cells^38^. Thus, the highly abundant γc cytokines present in adult lungs may fuel local antiviral effects further explaining the observed age differences in MCMV control. In addition, IL-33, IFN-α, and IL-27 were more abundant in adult than in neonatal mice. IL-33 promotes antiviral CD8 T cell immunity^39^ and IFN-α itself is known as a signal 3 cytokine affecting T cell activation, proliferation, and survival^40^. IL-27 has been shown to exert synergistic effects on TCR-dependent T cell proliferation and IL-27 receptor deficiency led to impaired expansion of potent anti-EBV effector cytotoxic CD8 T cells in humans^41^. Chronically MCMV infected *IL-27ra^-/-^* mice exhibit no prominent phenotype in the antiviral CD8 T cell pool but the role of IL-27 in the early T cell priming phase has not been studied^42^. Together, several cytokines that were present in higher concentrations in adults are involved in driving protective antiviral CTL immunity suggesting that a deficiency of these cytokines in the early-life T cell priming phase caused ineffective CD8 T cell responses.

T cells were activated in neonatal lungs indicating that the presentation of MCMV antigens is functional in early life. Indeed, in a previous study we found that NIFs contain various types of antigen-presenting cells (APC) and serve as priming sites for MCMV-specific CD8 T cells^3^. Recently, a subset of CD103^+^ APCs has been proposed to dampen CD8 T cell responses leading to impaired pulmonary immunity in the first two weeks of life in neonates^43^. Thus, besides the lack of several signal 3 cytokines, direct APC – T cell interaction may further interfere with early-life generation of antiviral CTLs.

In contrast to the CD8 T cell response, CD4 T cells did not reveal an age-related defect in effector differentiation. Rather, CD4 T cells primed in early life acquired a Th1-like phenotype with an even higher cytotoxicity score and increased expression of *Ifng*. Accordingly, we found more IFN-γ in MCMV-infected lungs in neonates than adults indicating that this cytokine counterbalances the absence of CTLs. This matches a study where T cell-produced IFN-γ was required to contain MCMV lung infection in adults^17^. It is well-described that CD4 T cells are involved in CMV control by promoting antibody production and direct antiviral effects in tissue^44,45^. However, their contribution appears minor to that of CD8 T cells^46^ and in this model they could not compensate the observed defect in CD8 T cell effector response. However, it is intriguing that the early-life T cell priming machinery selectively interferes with the generation of CTLs but not CD4 Th1 immunity and this needs further investigation.

We found higher frequencies of NK and γδ T cells in HCMV^+^ newborns (Fig S5A) and adaptive-like expansion of NKG2C^+^ NK cells and vδ1^+^ T cells occurs upon infection^47–51^. These findings meet their equivalent in mice where NK cells can kill infected cells via recognition of the MCMV-encoded m157 protein by the activating receptor Ly49H leading to subsequent expansion of the Ly49H^+^ NK cell pool^52^. Importantly, NK cells are present in neonatal mice at birth and present a relatively stable population of immune cells if compared to T cells (Fig1E+F). In line, γδ T cells are present in the early life and exhibit antiviral function to MCMV and HCMV^53^. Together, there is a cellular antiviral innate immune response and its contribution in supplementing αβ T cell immunity in the early-life phase needs further investigation.

T cells are important for control of CMV and effector CD8 T cells are present in newborns after HCMV infection^11^. This study argues that a detailed profiling of CMV-responsive CD8 T cells is required to allow correlation with protective function. Neonatal mice lack αβ T cells at birth and similarly, there are no αβ T cells in humans before the end of the 1^st^ trimester of gestation^54^. The number of T cells gradually increases during gestation supporting the hypothesis that the inverse correlation of gestational age at HCMV infection and risk of symptomatic cCMV^55–57^ is linked to the number of T cells present in the unborn fetus. In addition to that, the CDR3 regions of TCRs exhibit lower variations due to the low expression of TdT in early life^58,59^. Thus, a low number of T cells that can respond to CMV antigens together with altered priming conditions leads to an ineffective antiviral T cell response and contributes to the high susceptibility to CMV early in life.

Future studies need to address which factors potentiate the imbalance of early-life CD8 T cell immunity to understand why most are asymptomatic and some develop CMV disease. In this context, the *in situ* presence of signal 3 cytokines and phenotype of APCs during human αβ T cell priming need further investigation. Assessment of early-life human CD8 T cell cytotoxicity fitness using functional assays is required to confirm the findings from the animal model. Finally, early-life antiviral innate lymphocytes need to be taken into account to form a clearer picture of the emergence of cCMV disease.

## Limitations of the study

The results of this study imply that an insufficient neonatal T cell priming impairs differentiation into antiviral CTLs. The low number of T cells in the early life limit the possibility to study their antiviral effects after differentiation under adult T cell priming conditions. Additionally, in this study we investigated T cell immunity in MCMV-infected lungs and although we think this is an important site of infection one has to be careful with extrapolating the findings to other organs. Moreover, we had access to human peripheral blood only and could not confirm our findings in human lung T cells.

## Acknowledgments

We thank Martin Messerle for providing MCMV-2DR, Romy Hackbusch for the immune phenotyping of the infant samples, Kati Tillack for helping with the analysis, and Hans-Willi Mittrücker for critically reading of the manuscript and discussion. We thank the NIH Tetramer facility for providing M25 MHC- II tetramers. Data acquisition was performed using instruments of the University Medical Center Hamburg-Eppendorf (UKE) Microscopy Imaging Facility and the UKE FACS Sorting Core Unit.

This work was supported by the Deutsche Forschungsgemeinschaft (STA 1549/2-1, BR 1730/7-1 (KFO296), STA 1549/1-2, and BR 1730/9-1), the *Stiftung für Pathobiochemie und Molekulare Diagnostik* (German Society of Clinical Chemistry and Laboratory Medicine), German Center of Infection Research (DZIF 07.001-Stahl), the *Werner Otto Stiftung*, and the Medical Faculty of the University of Hamburg to FRS.

## Author Contributions

Conceptualization, F.R.S. Methodology, F.R.S. Investigation, L.F.B., S.T., E.O., A.P., D.I., R.B., A.G., E.T., and F.R.S.

Writing – Original Draft, F.R.S. and L.F.B. Writing –Review & Editing, F.R.S., L.F.B., E.T., P.A., L.G., and W.B. Funding Acquisition, F.R.S. and W.B. Resources, R.A., P.A., A.D., A.G., and W.B. Data Curation, S.V., L.G. Supervision, F.R.S. and W.B.

## Table Legends

Table S1. Differentially expressed genes of Th1 CD4 cells isolated from MCMV-infected adult and neonatal mice. Related to Figure 3.

Table S2. Differentially expressed genes of effector CD8 cells isolated from MCMV-infected adult and neonatal mice. Related to Figure 4.

Table S3. (B) Immune phenotyping of HCMV-exposed neonates. Related to Figure S5. Statistical differences between infected and non-infected newborns were calculated with a Mann-Whitney U test with a false discovery rate.

## Lead contact

Further information and requests for resources and reagents should be directed to and will be fulfilled by lead contact, Felix Rolf Stahl.

## Materials availability

MCMV-4DR was generated in this study and is available from the lead contact upon request.

## Data and code availability

Single-cell RNA-seq data has been deposited at ENA.

Any additional information required to reanalyse the data reported in this paper is available from the lead contact upon request.

## EXPERIMENTAL MODEL AND STUDY PARTICIPANT DETAILS

### Animals

C57BL/6J were purchased from Charles River Laboratories and bred in individually ventilated cages under specific pathogen-free conditions at the animal facility of the University Medical Center Hamburg-Eppendorf. β-actin-eGFP^60^, β-actin-eCFP^61^, *Rag2^-/-^IL2rg^-/^*^62,63^*^-^* and TCR transgenic OT-I^64^ and OT-II^65^ mice were all kept from a C57BL/6 background. OT-I mice were crossed with β-actin ECFP to generate OT-ICFP mice and OT-II mice with β-actin EGFP to generate OT-IIGFP mice. Both male and female mice were used for this study. Neonatal mice within 24h of birth and adult mice from 6 to 36 weeks of age were used. Experiments were performed according to the guidelines of the FELASA and Society of Laboratory Animals (GV-SOLAS) and approved by local authorities (Behörde für Gesundheit und Verbraucherschutz, Amt für Verbraucherscutz, Freie und Hansestadt Hamburg, reference numbers 06/16, 45/19 and 04/20).

### Human donors

Peripheral blood from HCMV-exposed neonates was collected within the first 15 days of life. Buffy coats from donors with known HCMV serology were obtained from the blood bank of the University Medical Center Hamburg-Eppendorf (UKE). The ethical permission for analysis of samples obtained from HCMV-exposed neonates was granted by the ethics committee of the Chamber of Physicians (Ärztekammer Hamburg) under the registration number (PV5689). At the time of study enrolment, parents provided written informed consent.

| ID | age (days), | gestational age at birth (days) | sex | CMV status | sample |
| --- | --- | --- | --- | --- | --- |
| CMV001 | 3 | 271 | male | negative | peripheral blood |
| CMV002 | 3 | 281 | female | negative | peripheral blood |
| CMV003 | 2 | 279 | male | positive | peripheral blood |
| CMV004 | 2 | 289 | female | negative | peripheral blood |
| CMV007 | 3 | 268 | female | negative | peripheral blood |
| CMV008 | 5 | 244 | female | negative | peripheral blood |
| CMV009 | 8 | 244 | female | negative | peripheral blood |
| CMV010 | 2 | 262 | male | negative | peripheral blood |
| CMV011 | 15 | 256 | male | negative | peripheral blood |
| CMV012 | 15 | 256 | male | negative | peripheral blood |
| CMV013 | 14 | 201 | female | negative | peripheral blood |
| CMV014 | 14 | 201 | female | negative | peripheral blood |
| CMV016 | 4 | 271 | male | negative | peripheral blood |
| CMV017 | 2 | 277 | female | negative | peripheral blood |
| CMV018 | 3 | 274 | male | positive | peripheral blood |
| CMV019 | 2 | 261 | female | negative | peripheral blood |
| CMV020 | 3 | 276 | female | negative | peripheral blood |
| CMV021 | 4 | 270 | female | negative | peripheral blood |
| CMV023 | 2 | 284 | male | negative | peripheral blood |
| CMV024 | 2 | 267 | female | negative | peripheral blood |
| CMV025 | 2 | 275 | male | negative | peripheral blood |
| CMV026 | 2 | 279 | male | positive | peripheral blood |
| CMV027 | 4 | 274 | male | positive | peripheral blood |

| ID | age (years) | sex | CMV status | sample |
| --- | --- | --- | --- | --- |
| BC02_22 | 71 | male | negative | frozen buffy coats |
| BC03_22 | 68 | male | negative | frozen buffy coats |
| BC21_21 | 58 | female | negative | frozen buffy coats |
| BC68_21 | 58 | female | negative | frozen buffy coats |
| BC74_21 | 63 | male | negative | frozen buffy coats |
| BC02_21 | 57 | male | positive | frozen buffy coats |
| BC25_21 | 63 | female | positive | frozen buffy coats |
| BC33_20 | 60 | male | positive | frozen buffy coats |
| BC67_21 | 71 | female | positive | frozen buffy coats |
| BC71_21 | 66 | male | positive | frozen buffy coats |

### Viruses

Infection experiments were performed with MCMV-3DR^66^ if not assigned differently. This recombinant consists of an *mCherry*-P2A-*Gaussia luciferase* insertion within the m157 ORF^67^ and additionally carries a sequence within the m164 ORF encoding the SIINFEKL peptide^68^. A previously described mutation with the m129 ORF has been repaired allowing full-length expression of MCMV-encoded chemokine 2 (MCK2) ^69^. MCMV-4DR is a modification of MCMV-3DR, where the sequence for a fusion protein made up of the first 118 residues of the human transferrin receptor linked to residues 299-385 from chicken ovalbumin was inserted into the *m128* ORF under control of the human major immediate early promotor (Fig. 2J). MCMV-2DR^70^ does not encode for ovalbumin peptides. All recombinants were derived from the pSM3fr Smith strain and modified by bacterial artificial chromosome mutagenesis using the *en passant* protocol^71^, propagated and titrated with 10.1 and M2-10B4 fibroblasts, respectively.

## METHODS DETAILS

### Infections

Neonatal mice were infected intratracheally within the first day of life (post-natal day 0) with 10^4^ PFU MCMV diluted in 10 µL PBS. Adult (6-36 weeks) mice were infected with 2x10^5^ PFU (10^6^ for experiments depicted in Figure 1) intranasally after anesthesia (100 mg/kg body weight ketamine and 5mg/kg body weight xylazine, intraperitoneal).

### Adoptive T cell transfers

Polyclonal T cells were isolated from spleens and lymph nodes by negative selection using a Pan T Cell Isolation Kit II (Miltenyi Biotec) according to the manufacturer’s guidelines. Naïve CD44^-^ T cells were isolated by magnetically depleting CD44^+^ cells using CD44 Microbeads (Miltenyi Biotec). Untouched OT-I and OT-II cells were isolated using a CD8a^+^ and CD4^+^ T cell Isolation Kit (Miltenyi Biotec), respectively. Stainings with Cell Proliferation Dye eFluor® 450 or 670 (eBioscience) were performed according to the manufacturer instructions.

CD44^+^ T cells were isolated from lungs of MCMV-infected *Wt* adult mice 7 dpi by positive selection of CD45^+^ cells with CD45 Microbeads (Miltenyi Biotec) followed by FACS cell sorting for CD3^+^CD44^+^ T cells. Cells were injected intraperitoneally at PND 0.

### Leukocyte isolation from lungs

Leukocytes were isolated after lung digestion using a Lung Dissociation Kit (Miltenyi Biotec) according to the manufacturer’s guidelines. Cells were then isolated via magnetic isolation or cell sorting as specified in the experiments.

### Histology

Organs were fixed in 2% PFA containing 30% sucrose, embedded in Tissue-Tek® O.C.T. medium and kept at -20°C until further processing. Cryosections of 7µm (slide scanner) or 10µm thickness (confocal microscopy) were stained after Fc receptor blocking with CD3-Alex Fluor 647 (17A2), B220-PE (RA3- 6B2), CD45-APC or CD45-Alexa Fluor 750 (30-F11) and DAPI. Images were acquired with a ZEISS Axioscan 7 Microscope Slide Scanner or with a Leica TCS SP8 confocal microscope. Images were processed with ZEN 2.6 or LAS AF Lite 4.0, respectively.

### Quantification of histology data

Infected cells and leukocytes from lungs were identified based on fluorophore expression (MCMV mCherry, OT-ICFP and OT-IIGFP), antibody or DAPI staining, and manually counted. In the adoptive transfers of OT-ICFP and OT-IIGFP T cells three to four NIFs per animal were analysed. In the remaining experiments, one whole lung section per animal was analysed. NIFs were identified based on preferential accumulation of CD45^+^ cells and/or nuclei, manually defined and areas calculated with ZEN 2.6.

### Flow cytometry, intracellular staining and cell sorting of mouse samples

Lung single cell suspensions were prepared as described above. Spleens and lymph nodes were mashed through 70µm cell strainers to obtain single cell suspensions and blood leukocytes were obtained after erythrocyte lysis for 15 min at room temperature. Cell surface stainings were then performed after blocking of Fc receptors with CD3-FITC, CD3-Brilliant Violet 711, CD3-Alexa Fluor 647 (17A2), CD3-PE (REA641)), CD4-PE-Vio770, CD4-PerCP (REA604), CD8b-APC (YTS156.7.7), CD8b-APC- Vio770 (REA793), CD44-APC, CD44-PE, CD44-Pacific Blue (IM7), CD44-PE-Vio770 (REA664), CD45-APC (30-F11), CD45R/B220-FITC (REA755), CD62L-Alexa Fluor 488 (MEL-14), CXCR6-Brilliant Violet 711 (SA051D1), KLRG1-APC-Cy7(2F1/KLRG1), NK1.1-APC (PK136), TCRvα2-PE (B20.1), TCRβ-PerCP-Vio700 (REA318) antibodies, M38-, M45-, m139-PE tetramers (provided by Ramon Arens) or M25-PE tetramer (NIH tetramer facility). For intracellular stainings, cells were fixed for 30 min after surface staining and incubated with CCL5-PE (2E9/CCL5) in permeabilization buffer. Single cell suspensions were measured within one day on a BD FACSCanto™ II or BD FACS LSRFortessa™ flow cytometer. Isolation by cell sorting was performed on a BD FACSAria™ Fusion.

### Flow cytometry human samples

For immune phenotyping infant blood samples, 50µl of whole blood were incubated with combinations of the following fluorochrome-conjugated antibodies for 30 min at room temperature: the “T-regulatory” panel contained anti-CD3 BV510 (clone: OKT3), anti-CD4 AF700 (clone: OKT4), anti- CD8 BV605 (RPA-T8), anti-CD25 BV421 (clone: BC 96), anti-CD31 APC-Cy7 (clone: WM59), anti-CD39 PE-Cy7 (A1), anti-CD45RA PE-Dazzle (clone: HI100), anti-CD73 PE (clone: AD2), anti-CD127 BV650 (clone: WM59), anti-HLA-DR BV711 (clone: L243), and anti-CCR4 PerCP-Cy-5.5 (clone: TG6); the “T-invariant” panel contained anti-CD3 BV510 (clone: OKT3), anti-CD4 PE-Dazzle (clone: RPA-T4), anti-CD8 AF700 (clone: HIT8a), anti-CD25 BV421 (clone: BC 96), anti-CD27 BV650 (clone: O323), anti-CD45RO BV785 (clone: UCHL1), anti-CD69 APC-Cy7 (clone: FN50), anti-CD161 BV605 (clone: HP-3G10), anti- CCR6 PerCP-Cy-5.5 (clone: G034E3), anti-CCR7 BV711 (clone: G043H7), anti-TCR γδ PE-Cy-7 (clone: 11F2), anti-Vα7.2 APC (clone: 3C10), anti-Vδ1 FITC (clone: TS-1), anti-Vδ2 APC (123R3) and anti-Vδ9 FITC (IMMU 360); finally, the “T-effector” panel contained anti-CD3 BV785 (clone: OKT3), anti-CD4 APC-Cy-7 (clone: RPA-T4), anti-CD8 BV510 (RPA-T8), anti-CD25 PE (clone: 2A3), anti-CD28 PE-Cy7 (clone: CD28.2), anti-CD38 AF700 (clone: HIT2), anti-CD45RA PE-Dazzle (clone: HI100), anti-CD57 FITC (clone: HCD57), anti-CD95 BV421 (clone: DX2), anti-CD127 BV650 (clone: WM59), anti-CCR7 APC (clone: G043H7) and anti-HLA-DR BV711 (clone: L243). After staining, 1ml of BD lysing solution was added the tube for lysis of erythrocytes, and stained cells were washed in PBS and resuspended in FACS buffer. Flow cytometric analysis was performed on an LSR Fortessa (BD Biosciences). Prior to analysis, a spillover spreading matrix was produced, PMT voltages were optimized and a compensation matrix was calculated. The FlowJo™ Software version 10.8.1 (FlowJo, LLC, Ashland, USA) was used for manual analysis, using a gating strategy similar to Mohme et al. ^72^ and Sibbertsen et al. ^73^.

### Cytokine multiplex assay

Mouse lungs were collected at 5 dpi after perfusion with PBS, weighed, and lysed with a TissueLyser II in 200µL PBS. Supernatants were frozen at -80°C until further processing. Supernatants were thawed on ice and processed with a LEGENDplex™ (Biolegend) custom made panel targeting IFN-α, IFN-β, IFN- γ, IL-10, IL-12p70, IL-15, IL-18, IL-1β, IL-2, IL-27, IL-33, IL-6, IL-7 and TGF-β1 (free active) according to the manufacturer’s guidelines. Samples were then measured on a BD FACSCanto™ II. Data analysis was performed on the LEGENDplex™ Data Analysis Software Suite (Biolegend).

### Luciferase

Organs were homogenized with a TissueLyser II in phosphate buffered saline, supernatants were measured in duplicates for *Gaussia luciferase* activity after automatic addition of Coelenterazine (Synchem) with a Centro LB 960 XS3 (Berthold). Luciferase values from non-infected animals were used as controls and/or to determine organ-specific limit of detection.

### scRNA

Adult *Wt* mice were infected with 2x10^5^ PFU MCMV-3DR. Naïve CD44^-^ T cells were isolated from age-matched eGFP mice using a combination of a magnetic cells sorting Pan T cell kit (Miltenyi Biotec) and subsequent CD44^+^ depletion as described above. After isolation, 10^7^ eGFP naïve CD44^-^ T cells were adoptively transferred into PND 0 neonates which were simultaneously infected with 10^4^ PFU MCMV- 3DR. At 7dpi, adult and neonatal mouse lungs were collected after perfusion with PBS and single cell suspensions were generated as described above. Lung T cells were then isolated by positive selection of T cells using CD3ε Microbeads (Miltenyi Biotec). Isolated cells were then stained with a CD3 antibody and FACS-sorted for CD3^+^ cells (adults) or CD3^+^eGFP^+^ cells (neonates). Single-cell RNA-seq libraries were constructed using a Chromium Single Cell 5’ Kit (v2) according to the manufacturer’s instructions.

### Single cell RNA-seq data analysis

#### Data processing

Raw FASTQ files were generated from Illumina sequencer’s base call files (BCLs) using the 10x Genomics Cell Ranger (version 4.0.0) ^74^ mkfastq pipeline. Count matrices of valid barcodes and feature/gene expression were generated by the Cell ranger count pipeline using mouse reference genome GRCm38 and GENCODE gene annotation version M23 provided by 10x Genomics. Custom GFP genome and annotation (using “MT-“ as a prefix) were added to the mouse genome, as well as to the mouse annotations, respectively.

#### Creating single cell objects and filtering

Filtered count matrices generated above for each sample were taken as input in Read10x function in R package Seurat (v4.2.0), and used for further downstream analysis. Following filters were used to create a scRNA-seq object using CreateSeuratObject function for each sample: 1) genes were removed if detected in less than 10 cells; 2) cells with a total number of genes below 200 were removed; 3) cells with a total number of genes detected above the 99^th^ percentile were removed; 4) cells with a proportion of mitochondrial UMIs of more than 5% were removed; and 5) cells with total UMIs less than 1000 were removed.

Separate datasets were created of object created above for *Cd8a* positive cells and *Cd4* positive cells, and following strategies were performed in similar way for sub-clustering as the original integrated scRNA-seq object with all samples.

#### Normalization, integration and downstream analysis

Normalization and variance stabilization of each scRNA-seq object was performed using statistical modelling framework sctransform^75^ implemented in Seurat, and also removing confounding source of variation such as: 1) total UMIs in a cell, 2) mitochondrial mapping percentage, and 3) eGFP expression. Resulting objects for all samples were integrated step by step using Seurat functions in order: SelectIntegrationFeatures (with parameter nfeatures=3000); PrepSCTIntegration; FindIntegrationAnchors and IntegrateData. Dimension reduction of integrated object created was performed using RunPCA function, and used as input to perform non-linear dimension reduction by using UMAP (Uniform Manifold Approximation and Projection) method with function RunUMAP using 30 dimensions. Prior to identify clusters of cells, shared nearest neighbor (SNN) graph was constructed using FindNeighbors function, and use as input for clustering by Louvain algorithm implemented in “FindClusters” function.

Default assay of the integrated object was switched to RNA and normalised using NormalizeData function to plot gene expression values. Default assay was switched back to integrated, and FindConservedMarkers function using default parameters was used to identify marker genes for all clusters of an integrated object. Cluster type identification and re-naming was done manually based on expression of the genes described in the results section.

Heatmap of top 20 marker genes in order of log2FC was created using “DoHeatmap” function. Cytotoxicity scores were added with the AddModuleScore including *Fasl*, *Gzma*, *Gzmb*, *Gzmc*, *Gzmk*, *Gzmm*, *Tnf*, *Tnfsf10*, *Prf1* and *Ifng* as features^76,77^.

#### TCR repertoire analysis

FASTQ files generated above were used with Cell Ranger vdj pipeline to perform V(D)J transcripts assembly and calling paired clonotypes, using V(D)J mouse reference (vdj-GRCm38-alts-ensembl-4.0.0) provided by 10x Genomics. Filtered contig annotation files for each sample from the pipeline were used to combine with clonotypes file using R script^78^ to create TCR repertoire metric files for each sample containing information such as: cell barcode; TCR clonotype IDs; v_gene; j_gene and CDR3 sequence. Clonotype IDs output by the Cell Ranger vdj pipeline were used to define clonotype frequency if more than one cell were represented by it.

scRNA-seq and TCR repertoire analysis, plots and tables were generated in R version 4.1.1 with ggplot2 version 3.3.6^79^.

#### Pseudo-bulk RNA-seq analysis

Gene expression matrix was created using AggregateExpression function on counts slot of each scRNA-seq object. DESeq2^80^ version 1.32.0 was used for normalisation and perform pairwise differential expression analysis.

## QUANTIFICATION AND STATISTICAL ANALYSIS

Statistical analyses were performed with GraphPad Prism 8 using the statistical tests as indicated. Radar charts were created using the function radarchart from the package fmsb^81^. Heatmap creation and UMAP dimensionality reduction of human samples were performed in R^78^ using the function pheatmap from the package pheatmap^82^ with standard complete clustering method and the function umap from the package umap^83^ with default parameters, respectively. Only if the number of samples was below the default value of 15, parameter n_neighbors from function umap was decreased according to the number of samples.

