## Supplemental Figures for "Limited protection against early-life cytomegalovirus infection results from deficiency of cytotoxic CD8 T cells"

### Supplemental figure legends

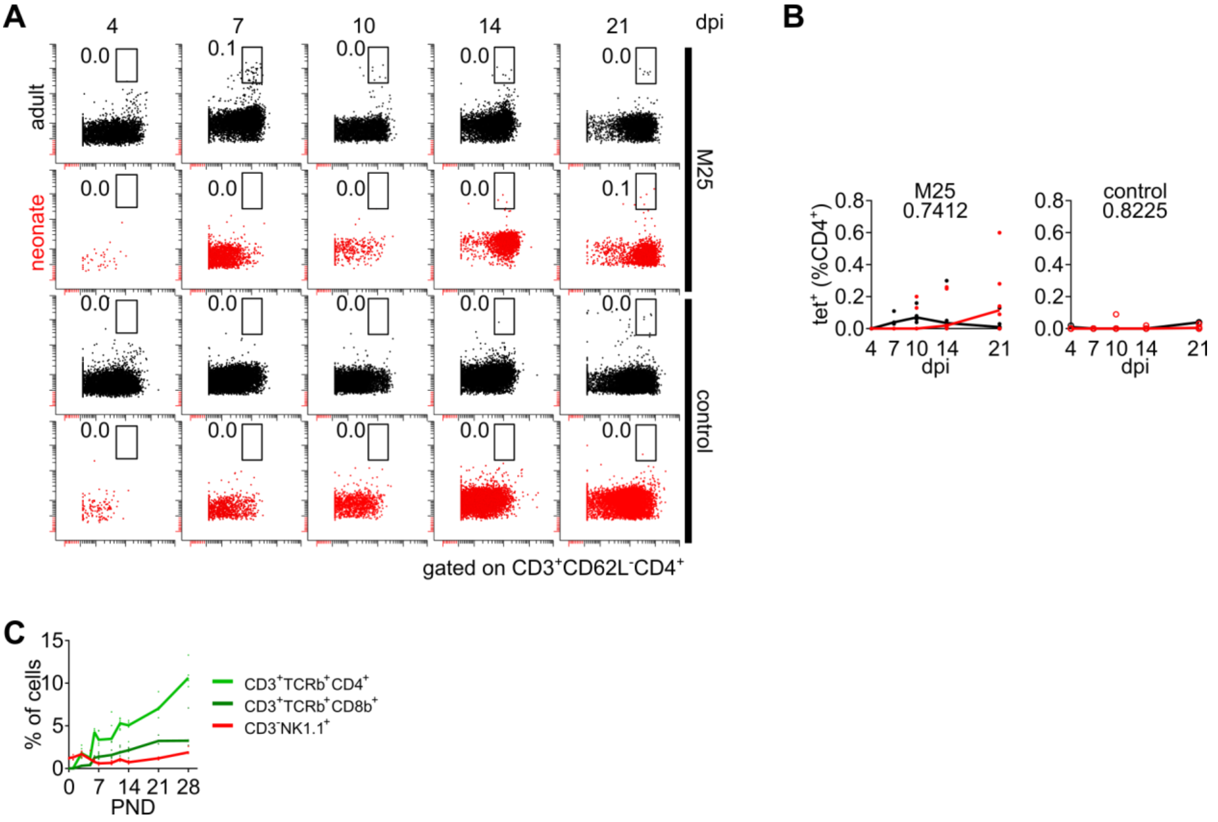


Fig S1. Detection of MCMV-specific CD4 T cells and accumulation of T and NK cells in spleens of non-infected mice. Related to Figure 1.

(A and B) (A) Representative flow cytometry plots and (B) pooled analysis of M25-specific CD4 T cells in the blood of infected and non-infected control mice.

(C) Accumulation of NK cells, CD4 and CD8 T cells in the spleen (data as depicted in Fig 1F, left panel but with adjusted y-axis scale).

Data display pooled results from 2 or more independent experiments (A and B, n=2-14 per time point, C, n=3-7 per time point). Lines in (B) indicate the median value. Numbers above each graph in (B) indicate the p value between adults and neonates of a 2-way ANOVA.

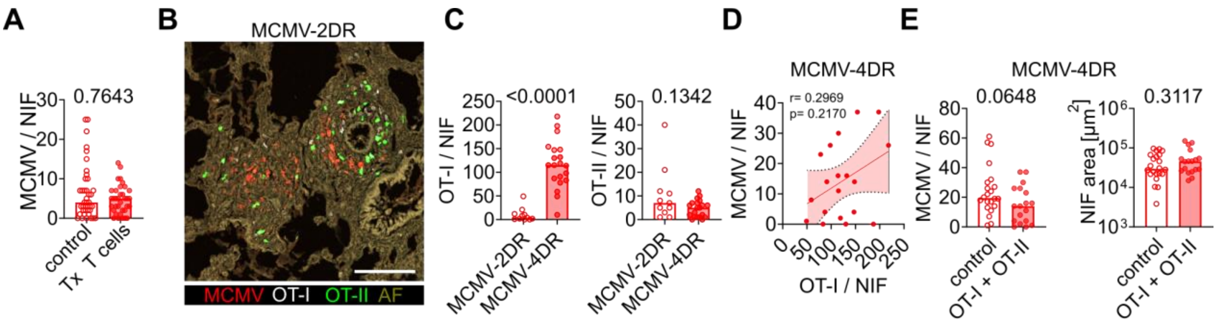


Fig S2. Adoptive transfer of adult naïve T cells into neonates is not protective against MCMV. Related to Figure 2.

(A) Quantitative analysis of lung immunohistology after adoptive transfer of adult polyclonal T cells. Related to Fig 2H.

(B) Representative immunohistology of a neonatal lung NIF after MCMV-2DR infection and adoptive transfer of OT-I and OT-II cells.

(C) Number of OT-I and OT-II cells in neonatal NIFs after MCMV-2DR or MCMV-4DR infection.

(D) Correlation of number of OT-I cells per NIF with number of MCMV-4DR-infected cells per NIF in neonates.

(E) NIF area and number of infected cells in NIFs of neonates infected with MCMV-4DR.

Data display pooled results from 3 or more independent experiments (A, n=7-11 per time point, B-D, n=3-4 per group). Numbers above each graph in (A), (C), and (E) indicate the p values of Mann-Whitney tests. (D) Nonparametric Spearman correlation r and two-tailed p value is provided.

A

| age | group | T cells | *Cd4*+ | *Cd8a*+ |
|---|---|---|---|---|
| adult | control | 1978 | 609 | 852 |
| adult | MCMV | 1559 | 538 | 836 |
| neonate | control | 1266 | 463 | 529 |
| neonate | MCMV | 1220 | 404 | 688 |
| | total | 6023 | 2014 | 2905 |

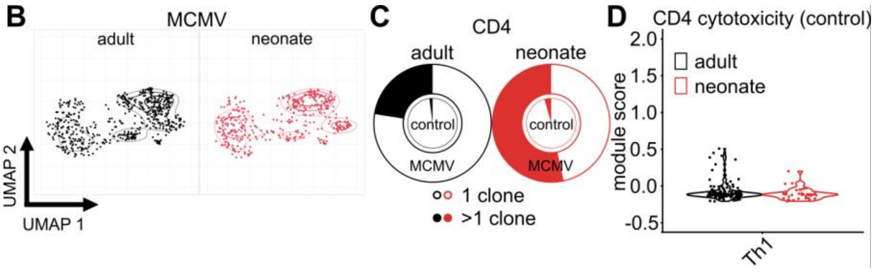


Fig S3. Single-cell RNA sequencing of T cells primed in adult and neonatal mice. Related to Figure 3.

(A) Absolute numbers of analysed T cells in each group.

(B) UMAP dimensionality reduction of CD4 T cells isolated from MCMV-infected animals.

(C) Relative distribution of clonal sizes of all CD4 T cells.

(D) Cytotoxicity module score of CD4 Th1 cells isolated from non-infected adults and neonates.

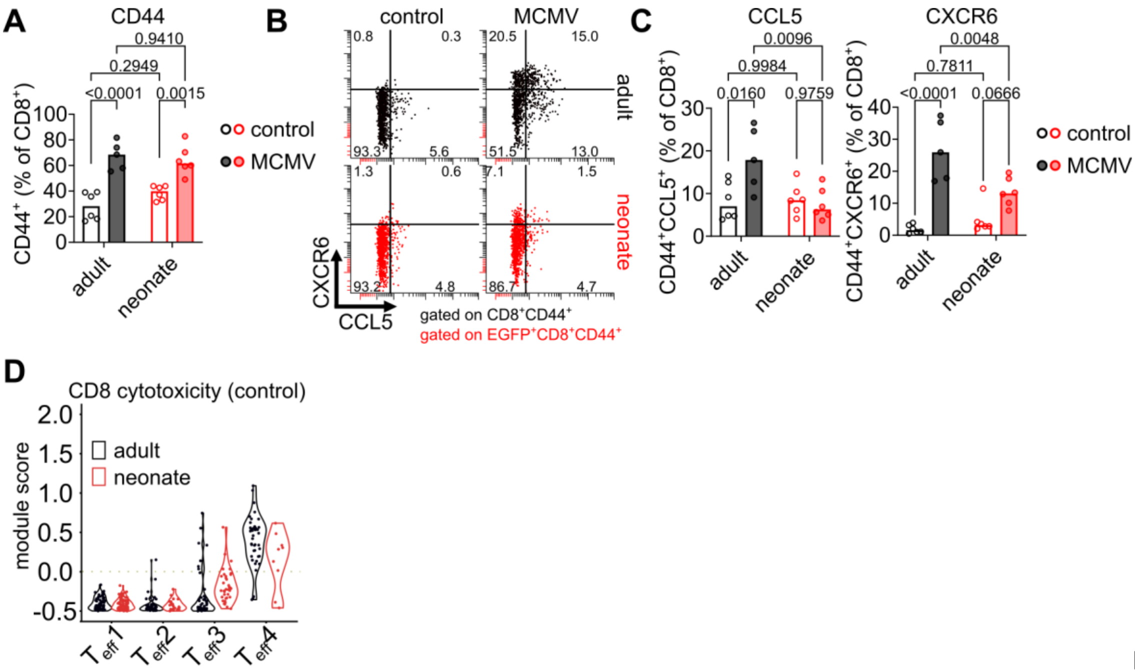


Fig S4.

Effector phenotype of CD8 T cells primed in adult and neonatal mice. Related to Figure 4.

(A) Analysis of CD44 protein expression by flow cytometry in adoptively transferred (eGFP$^+$) or endogenous CD8$^+$ T cells of neonates and adults, respectively.

(B and C) (B) Representative flow cytometry plot and (C) pooled analysis of CXCR6 and CCL5 expression in adoptively transferred (eGFP$^+$) or endogenous CD8$^+$CD44$^+$ T cells of neonates and adults, respectively.

(D) Cytotoxicity module score of effector CD8 T cells isolated from non-infected adults and neonates.

Data in A-C display pooled results from 2 independent experiments (n=5-6). Numbers above each graph in (A) and (C) indicate the p values of 2-way ANOVAs.

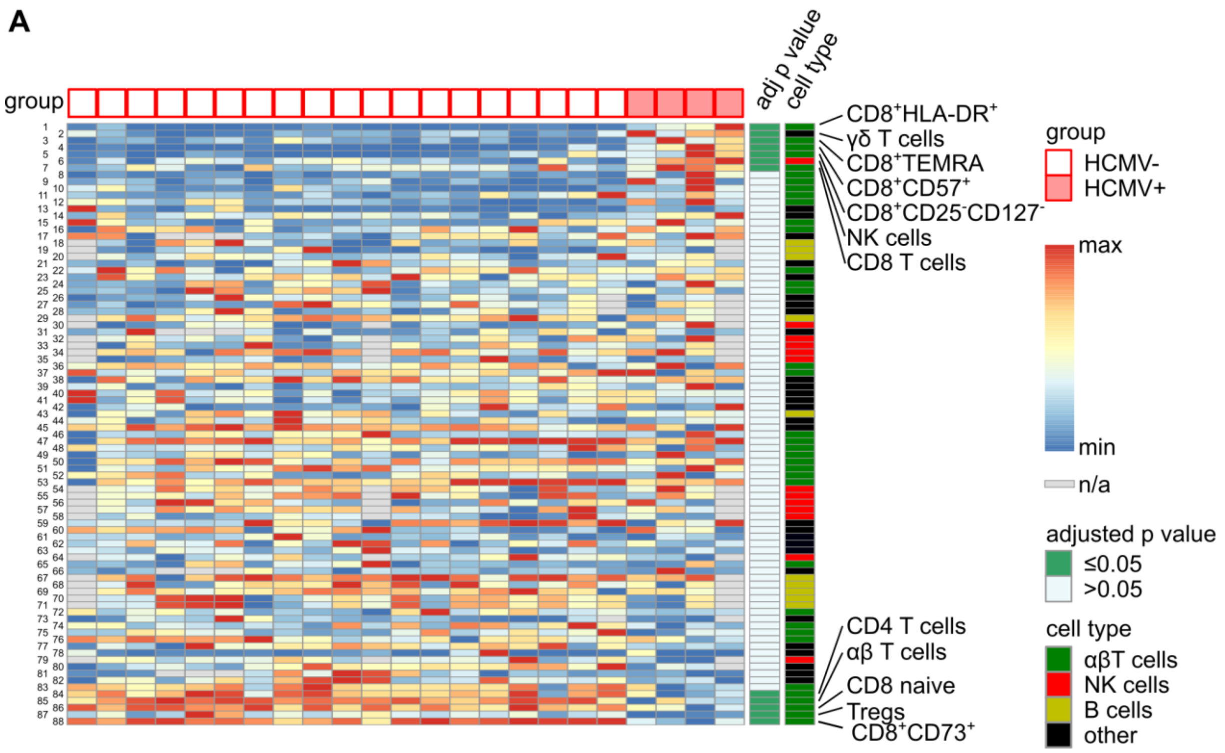


Fig S5. Immune phenotyping of HCMV-exposed neonates. Related to Figure 7.

(A) Heatmap showing relative frequencies of all peripheral blood populations measured in HCMV-exposed neonates. Numbers in row refer to cell populations as described in Table S3. Values are normalized to minimum and maximum values. Adjusted p-values for comparison of HCMV$^-$ and HCMV$^+$ individuals were calculated using Mann-Whitney U tests with false discovery rate correction.
